# Longitudinal *in vivo* micro-CT-based approach allows spatio-temporal characterization of fracture healing patterns and assessment of biomaterials in mouse femur defect models

**DOI:** 10.1101/2020.10.02.324061

**Authors:** Esther Wehrle, Duncan C Tourolle né Betts, Gisela A Kuhn, Erica Floreani, Malavika H Nambiar, Bryant J Schroeder, Sandra Hofmann, Ralph Müller

## Abstract

Thorough preclinical evaluation of functionalized biomaterials for treatment of large bone defects is essential prior to clinical application. Using *in vivo* micro-computed tomography (micro-CT) and mouse femoral defect models with different defect sizes, we were able to detect spatio-temporal healing patterns indicative of physiological and impaired healing in three defect sub-volumes and the adjacent cortex. The time-lapsed *in vivo* micro-CT-based approach was then applied to evaluate the bone regeneration potential of functionalized biomaterials using collagen and BMP-2. Both collagen and BMP-2 treatment led to distinct changes in bone turnover in the different healing phases. Despite increased periosteal bone formation, 87.5% of the defects treated with collagen scaffolds resulted in non-unions. Additional BMP-2 application significantly accelerated the healing process and increased the union rate to 100%. This study further shows potential of time-lapsed *in vivo* micro-CT for capturing spatio-temporal deviations preceding non-union formation and how this can be prevented by application of functionalized biomaterials.

This study therefore supports the application of longitudinal *in vivo* micro-CT for discrimination of normal and disturbed healing patterns and for the spatio-temporal characterization of the bone regeneration capacity of functionalized biomaterials.

## Introduction

Regeneration and healing of large bone defects (e.g. caused by trauma, infection, tumor resection, congenital skeletal disorders) is a treatment challenge in orthopedic surgery with as much as 10-20% of patients experiencing delayed or non-unions ^1–3^. Recent advances in tissue engineering and material sciences (e.g. 3D-bioprinting) enabled the development of diverse biomaterials, which can be functionalized with biochemical factors (e.g. growth factors) and combined with cell therapeutic approaches ^3–5^. In order to facilitate the clinical application of these innovative approaches for the treatment of large bone defects, their bone regeneration capacity needs to be systematically and thoroughly characterized in preclinical studies ^6–8^. For this purpose, critical size defect models have been developed for load-bearing and non-load-bearing bones in small and large animals ^7,9–13^. So far, most of these studies focused their evaluation on end-point radiological and histological analysis ^14–16^. However, recent studies indicate that longitudinal non-invasive imaging could improve the evaluation of a biomaterial’s bone regeneration capacity, due to the ability to follow the regeneration process in the same animal over time, thereby also reducing animal numbers according to the 3R’s of animal welfare ^17,18^. Particularly, *in vivo* micro-CT was shown to be suitable for the assessment of bone tissue formation and mineralization after biomaterial application in critical size defect models ^17–19^. A further development is the consecutive registration of time-lapsed *in vivo* images ^20^. We recently developed a longitudinal *in vivo* micro-CT-based approach for healing-phase-specific monitoring of fracture repair in mouse femur defect models ^21^. Registration of consecutive scans using a branching scheme (bridged vs. unbridged defect) combined with a two/multi-threshold approach enabled the assessment of localized bone turnover and mineralization kinetics relevant for monitoring callus remodelling ^21,22^. Furthermore, we showed that longitudinal *in vivo* micro-CT imaging itself did not significantly affect callus formation and remodelling ^22^.

It is well accepted that longitudinal non-invasive imaging of the healing process is advantageous compared to cross-sectional study designs^22^. In order to apply *in vivo* micro-CT-based longitudinal monitoring approaches for the evaluation of functionalized biomaterials, it has to be assessed, whether these approaches allow for reliable discrimination between normal and impaired healing conditions (e.g. critical-sized defects). So far, preclinical studies were often not able to reliably capture the bone healing potential of biomaterials due to limitations in study design: (I) cross-sectional setup, (II) assessment not considering defect sub-volumes, (III) assessment not specific to the healing phases.

Therefore, this study assesses whether our recently developed time-lapsed micro-CT based monitoring approach is suitable for discrimination of healing patterns associated with physiological and impaired bone healing conditions using mouse femur defect models with different gap sizes. In a second step, we evaluated whether the time-lapsed *in vivo* micro-CT-based monitoring approach is suitable for profound characterization of the bone regeneration capacity of functionalized biomaterials using well characterized porous collagen scaffolds and bone morphogenetic protein (BMP-2) as gold standard

## Results

In Experiment 1, we assessed the potential of time-lapsed *in vivo* micro-CT for reliable discrimination between physiological and impaired fracture healing patterns. We compared the healing process (week 0-6) in mice receiving either a small (0.9mm, n=10) or large femur defect (2mm, n=8; Fig. 1; Supplementary Video 1). Via registration of consecutive micro-CT scans, structural and dynamic callus parameters were followed in three callus sub-volumes (defect center, DC; defect periphery, DP; cortical fragment periphery, FP) and the adjacent cortical fragments (FC) over time (Fig. 1, 2). In Experiment 2, we applied the same time-lapsed *in vivo* imaging approach to assess the bone regeneration capacity of collagen scaffolds (n=8) and collagen scaffolds+BMP-2 (n=8; Fig. 3, 4, Supplementary Video 2; for detailed study design see Supplementary Table S1).

**Figure 1.**
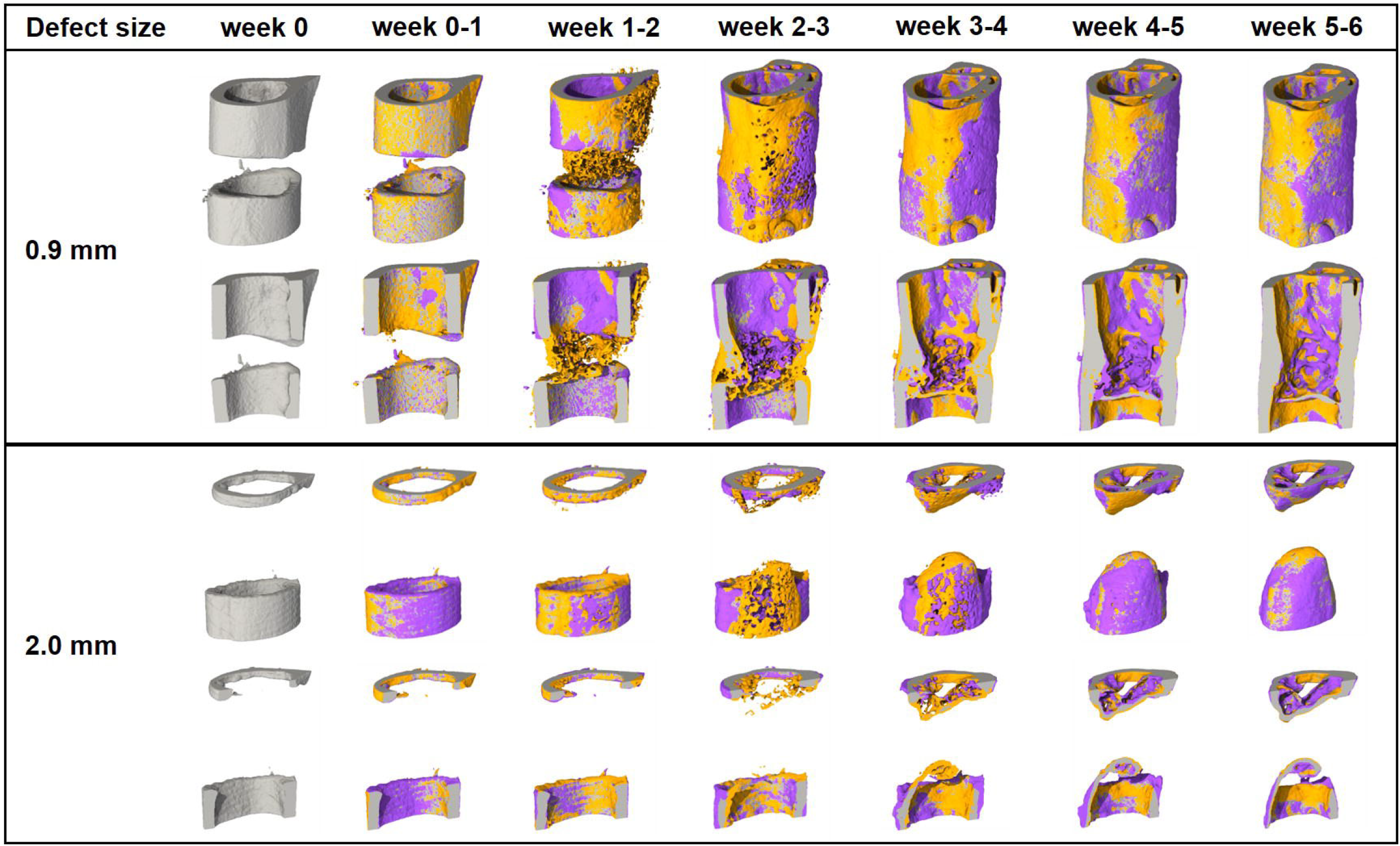
Representative images (threshold: 645 mg HA/cm^3^) of the defect region from animals of the 0.9mm group (top) and the 2.0mm group (bottom). Visualization of bone formation (orange) and resorption (blue) via registration of micro-CT scans from weeks 1-6 to weeks 0-5.

**Figure 2.**
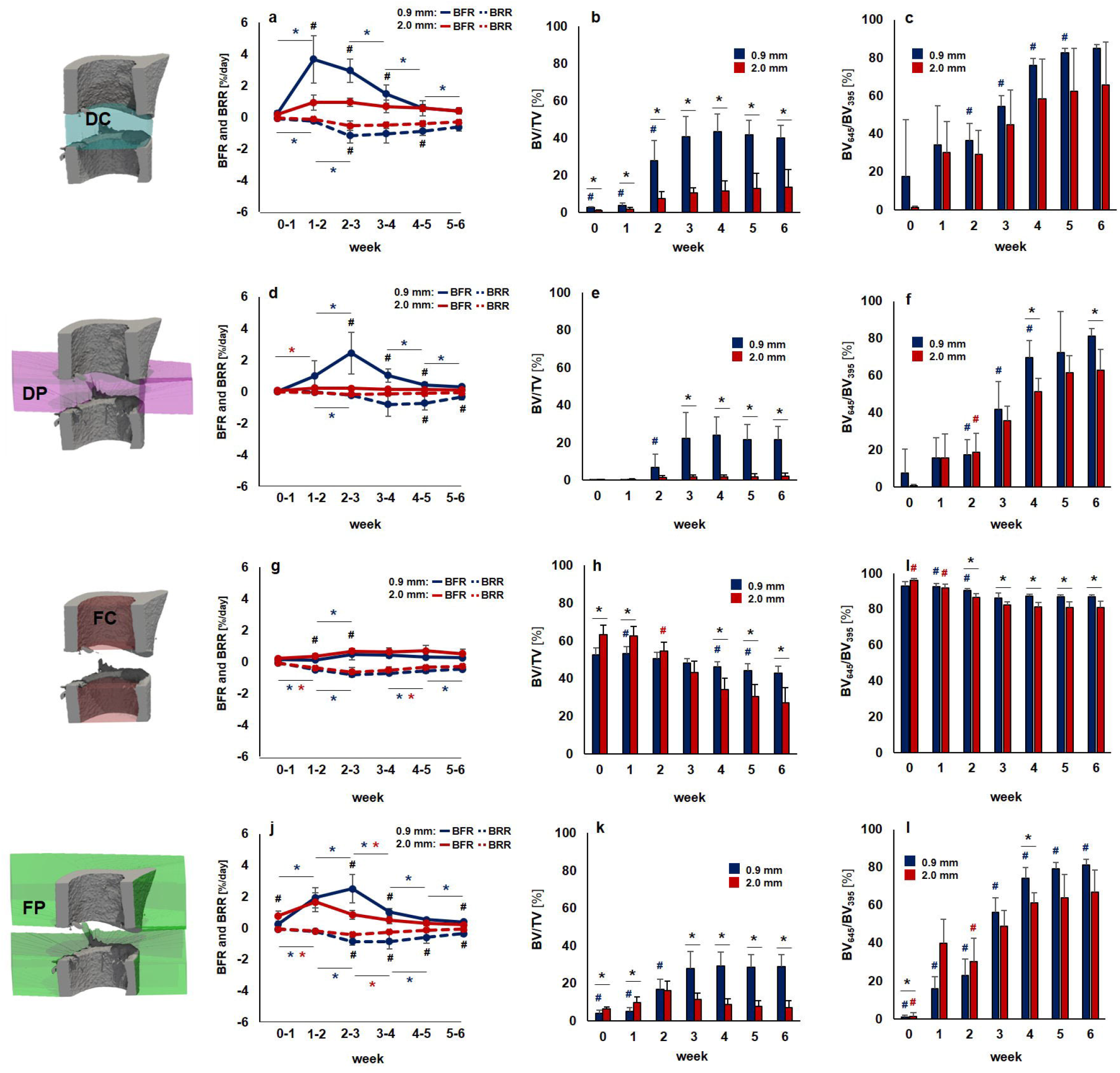
Micro-CT based evaluation of bone parameters in the 0.9mm group (blue) and the 2.0mm group (red) using different VOIS: defect center (DC; a-c), defect periphery (DP; d-f), cortical fragment center (FC; g-i), cortical fragment periphery (FP; j-l). a, d, g, j: Bone formation rate (solid line) and bone resorption rate (dashed line) given in percent per day. b, e, h, k: Bone volume (BV) normalized to TV (DC for DC and DP, FC for FC+FP). c, f, i, l: Degree of bone mineralization given as ratio of bone volume with a density ≥645 mg HA/cm^3^ to the total osseous volume (threshold ≥395 mg HA/cm^3^). n=7/10; a,d,g,j: * indicates p < 0.05 between consecutive weeks; ^#^indicates p < 0.05 between groups. b,c,e,f,h,I,k,l: * indicates p < 0.05 between groups; ^#^ indicates p < 0.05 between consecutive weeks.

**Figure 3.**
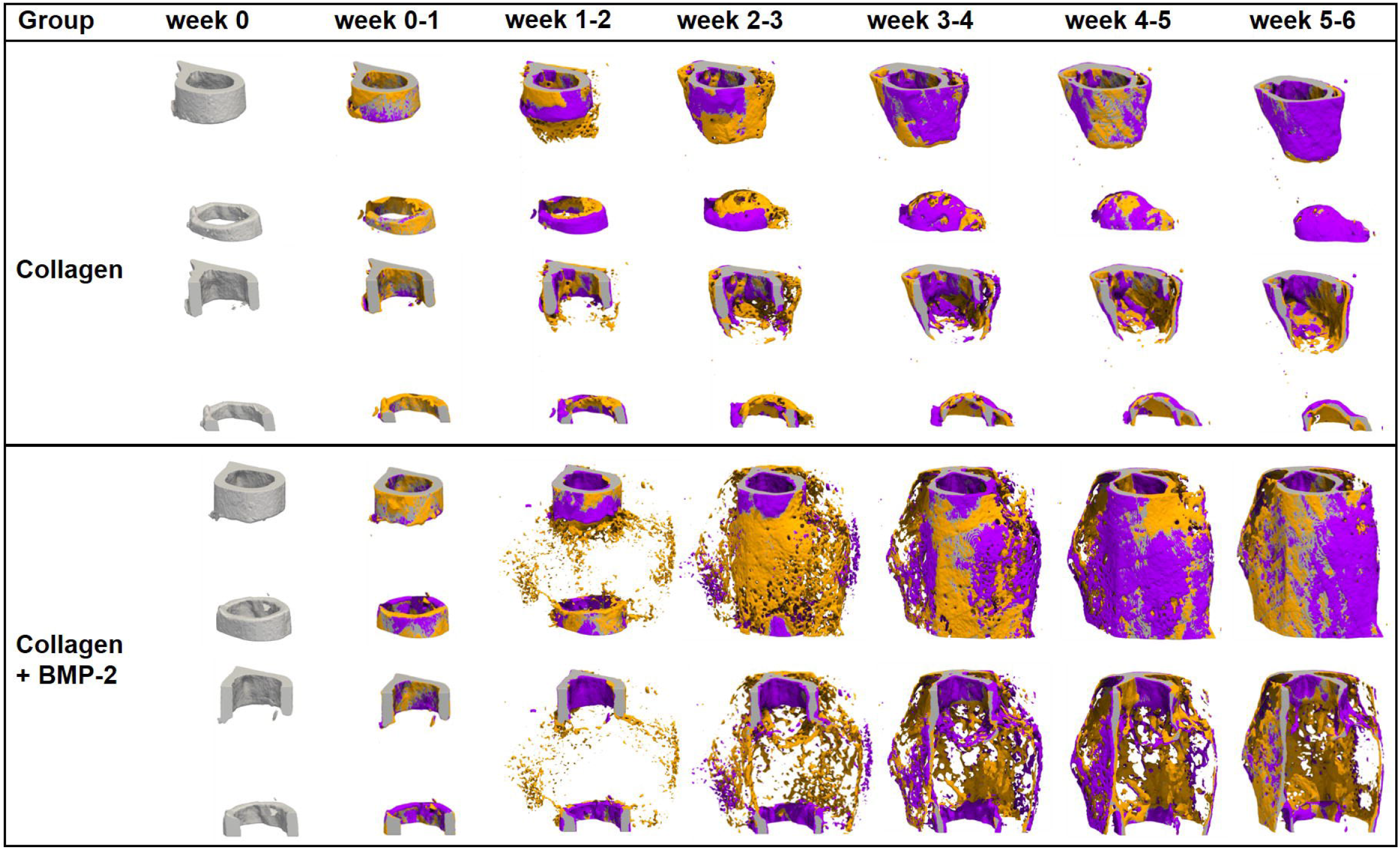
Representative images (threshold: 645 mg HA/cm^3^) of the defect region from animals of the collagen group (top) and the collagen+BMP-2 group (bottom). Visualization of bone formation (orange) and resorption (blue) via registration of micro-CT scans from weeks 1-6 to weeks 0-5.

**Figure 4.**
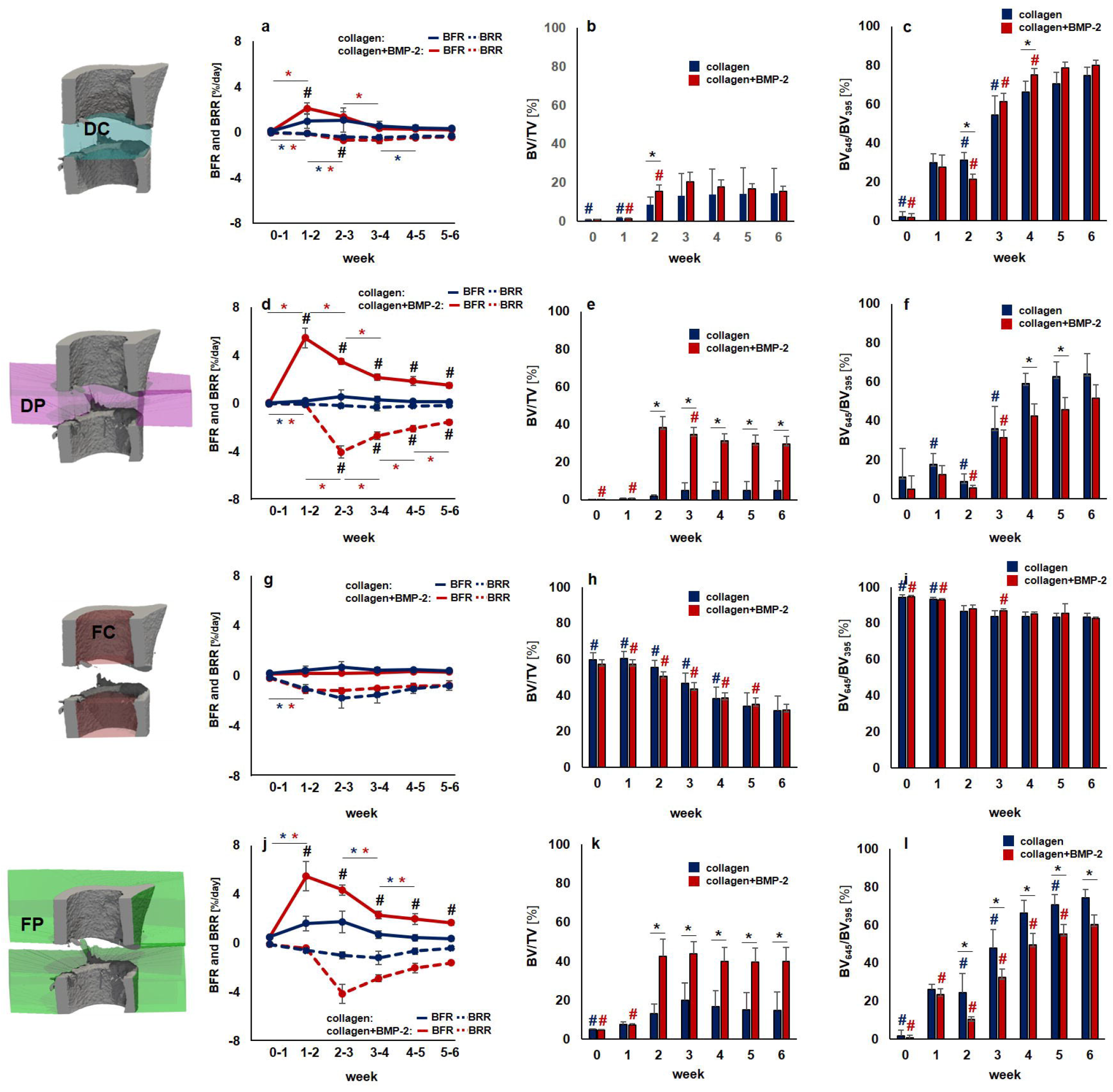
Micro-CT based evaluation of bone parameters in the collagen group (blue) and the collagen+BMP-2 group (red) using different VOIS: defect center (DC; a-c), defect periphery (DP; d-f), cortical fragment center (FC; g-i), cortical fragment periphery (FP; j-l). a, d, g, j: Bone formation rate (solid line) and bone resorption rate (dashed line) given in percent per day. b, e, h, k: Bone volume (BV) normalized to TV (DC for DC and DP, FC for FC+FP). c, f, i, l: Degree of bone mineralization given as ratio of bone volume with a density ≥645 mg HA/cm^3^ to the total osseous volume (threshold ≥395 mg HA/cm^3^). n=7/10; a,d,g,j: * indicates p < 0.05 between consecutive weeks; ^#^ indicates p < 0.05 between groups. b,c,e,f,h,I,k,l: * indicates p < 0.05 between groups; ^#^ indicates p < 0.05 between consecutive weeks.

### General physical observation

Post-operative monitoring was performed with a scoring system defined in license number 36/2014 (Kantonales Veterinäramt Zürich, Zurich, Switzerland), evaluating the following parameters: social behaviour, body position, motion, load bearing of operated limb, habitus, surgery wound. Scoring values: 0-4 (0=normal, 4=severely impaired, end-point criteria: scored 4 in one category or overall score ≥8). Monitoring schedule: evening of surgery day, day 1-3: morning and evening, day 4 until end of experiment: 3x/week, daily if scored ≥2 in one criterion. All mice recovered rapidly from surgery. However, one animal from the 0.9mm defect group only showed minor load bearing of the operated limb in the first post-operative week, which then gradually increased, reaching normal values after 2 weeks. The body weight did not significantly change during the healing period and did not differ between the groups in Experiment 1 and 2 (see Supplementary Fig. 1 and 2). Social interaction between mice and nesting behaviour did not differ from pre-surgical observations and were similar for animals from all groups.

### VOIs for evaluation by time-lapsed *in vivo* micro-CT

#### Experiment 1

In order to exclude bias in the further micro-CT analyses, we compared the size of the two central VOIs (DC, FC) used for normalization of CT parameters (depicted in Fig. 2) between groups. One animal from the 2.0mm group could not be included in the analysis due to incorrect registration caused by differences in leg alignment between micro-CT scans. The two central VOIs encompassed the following volume for the 0.9mm group (n=10) and the 2.0mm group (n=7) with significant group differences given in brackets: 1.64±0.19mm^3^ vs. 3.45±0.18mm^3^ for DC (p<0.0001), and 3.18±0.12mm^3^ vs. 0.94±0.30mm^3^ for FC (p<0.0001).

#### Experiment 2

The DC and FC VOIs encompassed the following volume for the collagen group (n=8) and the collagen+BMP-2 group (n=8): 3.09±0.22mm^3^ vs. 3.15±0.22mm^3^ for DC and 1.25±0.26mm^3^ vs. 1.47±0.19mm^3^ for FC. No significant group differences in VOI size were detected between the two groups.

### Longitudinal monitoring of fracture healing by time-lapsed *in vivo* micro-CT in non-critical and critical-sized femur defects (Experiment 1)

In both groups the repeated micro-CT scans (1x/week, Fig. 1, Supplementary Video 1) covered the period from the day of the defect surgery (d0) until post-operative week 6. In the 0.9mm group, distinct callus characteristics indicative of the different healing phases (inflammation, repair, remodelling) were seen in the three callus VOIS (DC, DP, FP; Fig. 1 and Fig. 2) as previously described in the same femur defect model ^22^. Specifically, a strong increase in bone formation with maximum values in week 1-2 (BFR: DC - 3.67±1.50%) and week 2-3 (BFR: DP - 2.44 ±1.31%, FP - 2.50±0.89%; Fig. 2) indicated the progression from the inflammation to the repair phase. This triggered bone resorption with maximum values seen in week 2-3 (BRR: DC - 1.15±0.48%) and week 3-4 (BRR: DP - 0.82±0.74%, FP - 0.86±0.48%) indicating progression to the remodelling phase. In all callus VOIs, a significant increase (p<0.0001) in bone volume was seen over time with maximum values observed in week 4 (BV/TV: DC - 44±9%, DP - 24±10%, FP - 29±8%). From week 2 onwards, the fraction of highly mineralized bone continuously increased in all callus VOIs until the study endpoint (week 6).

In the 2.0mm defect group, only a slight onset in bone formation was seen in the callus VOIs from week 0-1 to week 1-2 without characteristic peak values in the further healing process (Fig. 1 and 2). We could also not observe any significant gain in bone volume throughout the healing period, indicating an impaired and delayed healing pattern. Nevertheless, we saw a continuous increase in the fraction of highly mineralized bone in all callus VOIs, indicating no substantial disturbances of callus mineralization, despite the small callus volume. Comparison of both groups (0.9mm defect, 2.0mm defect) showed similar patterns in healing initiation with bone formation, first starting in the defect center (DC) and the cortical fragment periphery (FP) from week 0-1 to week 1-2. However, in the DC VOI the increase in bone formation (6x) was significantly lower (p=0.001) in the 2mm group compared to the 0.9mm group (19x) and the bone formation rates then remained stable at low values (≤0.95%/day) throughout the healing period associated with only little endosteal callus formation. In the FP VOI, the increase in bone formation from week 0-1 to week 1-2 was similar for both groups, but the 2.0mm defect group then showed a sudden decline (−40%) in bone formation (week 1-2 vs week 2-3) leading to premature cessation of periosteal callus formation. In the defect periphery (DP), hardly any callus formation (BFR ≤0.21%/day) was seen in the 2.0mm defect group throughout the healing period leading to significantly smaller callus dimensions compared to the 0.9mm group from week 3 until the study end (p=0.0001). With respect to bone resorption, no significant differences between the two defect groups were seen in the early healing period (week 0-1, week 1-2) in any of the callus sub-volumes. However, in the later healing period, bone resorption was significantly lower in the 2.0mm defect group compared to the 0.9mm defect group in DC (week 2-3: p=0.047, week 4-5: p=0.0064), FP (week 2-3: p= 0.0045, week 5-6: p=0.0048) and DP (week 4-5: p=0.0157, week 5-6: p=0.0177) VOIs, indicating impaired bone remodelling. By week 6, not only the bone volume, but also the fraction of highly mineralized bone was lower in all callus sub-volumes in the 2.0mm defect group compared to the 0.9mm defect group with significant differences (p=0.0383) being observed in the DP sub-volume (BV_645_/BV_395_: 2.0mm group - 63±11%, 0.9mm group - 82±4%).

To assess functional healing outcome, we particularly focused on the defect VOIs (DC+DP) which are most important for evaluating later healing time points during the remodelling phase of fracture healing. The previously observed differences in bone turnover, bone volume and mineralization between the two groups also affected cortical bridging. According to the standard clinical evaluation of X-rays, the number of bridged cortices per callus was evaluated in two perpendicular planes and animals with ≥3 bridged cortices were categorized as healed. Cortical bridging first occurred by week 3 in 70% of the animals in the 0.9mm defect group, whereas none of the animals in the 2.0mm group showed bridged cortices at this time point (Table 1). By week 6, 90% of the 0.9mm defects were categorized as healed with only 1 defect being classified as non-union. In contrast, in the 2.0mm defect group only 1 animal showed cortical bridging and 87.5% manifested as non-unions.

**Table 1.** Number of bridged cortices per callus evaluated weekly in two perpendicular planes and number of mice with successful fracture healing in the 0.9mm and the 2.0mm defect groups (≥ 3 bridged cortices, threshold 395 mg HA/cm^3^).

| Week | Group | Number of bridged cortices |  |  |  |  | Fracture healing outcome |  |
| --- | --- | --- | --- | --- | --- | --- | --- | --- |
|  |  | 0 | 1 | 2 | 3 | 4 | Not healed | Healed |
| 0 | 0.9mm | 10 | 0 | 0 | 0 | 0 | 10 | 0 |
|  | 2.0mm | 8 | 0 | 0 | 0 | 0 | 8 | 0 |
| 1 | 0.9mm | 10 | 0 | 0 | 0 | 0 | 10 | 0 |
|  | 2.0mm | 8 | 0 | 0 | 0 | 0 | 8 | 0 |
| 2 | 0.9mm | 7 | 3 | 0 | 0 | 0 | 10 | 0 |
|  | 2.0mm | 8 | 0 | 0 | 0 | 0 | 10 | 0 |
| 3 | 0.9mm | 1 | 0 | 2 | 6 | 1 | 3 | 7 |
|  | 2.0mm | 7 | 1 | 0 | 0 | 0 | 8 | 0 |
| 4 | 0.9mm | 1 | 0 | 1 | 5 | 3 | 2 | 8 |
|  | 2.0mm | 7 | 0 | 0 | 1 | 0 | 7 | 1 |
| 5 | 0.9mm | 1 | 0 | 0 | 4 | 5 | 1 | 9 |
|  | 2.0mm | 7 | 0 | 0 | 1 | 0 | 7 | 1 |
| 6 | 0.9mm | 1 | 0 | 0 | 4 | 5 | 1 | 9 |
|  | 2.0mm | 7 | 0 | 0 | 1 | 0 | 7 | 1 |

Bone turnover in the cortical fragments (FC) showed a similar pattern in both groups, without characteristic peak values in bone formation and resorption rates. Specifically, maximum bone formation rates were significantly lower (0.9mm group: −80%, p<0.0001, 2.0mm group: −58%, p=0.004) compared to those observed in the periosteal region (FP). Furthermore, a negative bone turnover was seen with maximum resorption rates (BRR in week 2-3; 0.79±0.15%) exceeding bone formation rates (BFR in week 2-3: 0.49±0.37%) in the 0.9mm group. This resulted in a decrease in bone volume over time (0.9mm group: −19%, 2.0mm group: −57%). Additionally, a significant 8% (0.9mm group, p=0.0017) and 11% (2.0mm group, p=0.0072) decrease in the fraction of highly mineralized bone was seen from week 1 to week 3 suggesting reorganization of the cortical bone adjacent to the forming and remodelling fracture callus.

### Longitudinal assessment of biomaterials by time-lapsed *in vivo* micro-CT in critical-sized femur defects (Experiment 2)

In the collagen group, no significant weekly changes in bone formation rate were seen in the central defect VOIs (DC, DP; Fig. 3 and 4; Supplementary Video 2). Bone resorption rates slightly but significantly increased during the early healing period (DC: 7.6x from week 0-1 to week 1-2, DP: 4.3x from week 0-1 to week 1-2) and then decreased (DC) or remained stable (DP) from week 2-3 to week 5-6. This led to only little callus formation in these central VOIs, as similarly seen in the 2mm defect group from Experiment 1, indicating an impaired endosteal fracture healing pattern associated with the application of the collagen scaffolds. In the periosteal VOI (FP), a significant 3.2x increase in bone formation rate was seen from week 0-1 to week 1-2, which remained stable during the subsequent week, before decreasing after week 2-3 reaching baseline values by week 5-6. No significant changes were seen in the bone resorption rate throughout the healing process. Compared to the 2mm defect group from Experiment 1, similar periosteal callus volumes were observed in the early healing period (BV/TV in week 2: collagen group: 13.08±4.88%, 2mm defect group: 15.99±5.09%). In the collagen group, the periosteal callus volume further increased (+53%) from week 2 to week 3, whereas it decreased (−30%) in the 2mm defect group during the same period, suggesting changes in periosteal callus formation potentially associated with the application of the collagen scaffolds.

In the collagen+BMP-2 group, a similar healing pattern compared to the collagen group was seen in the defect center (DC). Nevertheless, BMP-2 application led to a significantly 2.1x increased bone formation rate in week 1-2 compared to the collagen group, which was also associated with a transiently increased mineralized callus volume in this endosteal VOI in week 2. However, in week 6, both groups showed similar callus volumes and fraction of highly mineralized bone, indicating only a slight and transient BMP-2 associated effect on callus formation and remodelling in the defect center. In contrast, in DP and FP VOIs a completely different picture was seen: compared to the collagen group, a sudden 17x (DP) and 3.6x (FP) induction in BFR already from week 0-1 to week 1-2 was detected. This indicates that collagen+BMP-2 scaffolds are able to induce bone formation at defect locations distant to the cortical bone potentially allowing healing of large defects. The BMP-2 induced increase in the bone formation rate was higher but persisted shorter compared to the 0.9mm defect group with uncompromised healing in Experiment 1. This led to a significant 318x (DP) and 9.1x (FP) increase in bone volume by week 2 (DP: p<0.0001; FP: p=0.0002). The periosteal callus volume significantly (p<0.0001 for DP and FP) exceeded the values of the collagen group and was also higher compared to the 0.9mm group from Experiment 1. In both VOIs a significant 63x (DP) and 10x (FP) increase in bone resorption was seen from week 1-2 to week 2-3 (p<0.0001 for both VOIs), indicating early onset of callus remodelling. The bone resorption rates then remained at significantly higher levels compared to the collagen group and both groups from Experiment 1, indicating pronounced callus remodelling in the collagen+BMP-2 group. The fraction of highly mineralized tissue remained significantly lower in the collagen+BMP-2 group indicating that the remodelling process was still ongoing, whereas the bone healing process had finished in all other groups.

Despite the BMP-2 associated improved healing outcome, the healing pattern was completely different compared to the 0.9mm defect group with uncompromised healing. In the collagen+BMP-2 group bone formation mainly took place in the periosteal regions and the defect periphery, whereas in the 0.9mm defect group bone formation mainly took place in the defect center. The collagen group showed similar values compared to the 2.0mm defect group indicating impaired healing, suggesting that the collagen scaffold might have blocked endosteal callus formation.

In the FC VOI, we saw a similar pattern in all indices for the collagen and the collagen+BMP-2 group without any significant differences between groups. Specifically, the bone formation rate did not significantly change over time in both groups. Similar to the 2mm defect group from Experiment 1, a significant induction of bone resorption rate was seen from week 0-1 to week 1-2. From week 1 until the study end, the bone volume significantly (p<0.0001) decreased in both groups (collagen group: −48%, collagen+BMP-2 group: −44%). Furthermore, a significant decline in the fraction of highly mineralized bone was observed in both groups from week 0 to week 6 (collagen group: −9%, p=0.0002; collagen+BMP-2 group: −13%, p<0.0001), as similarly observed for the 2mm defect group in Experiment 1 (−16%, p=0.003).

Cortical bridging occurred by week 2 in 100% of the animals in the collagen+BMP-2 group, whereas no animal in the collagen group showed bridged cortices at this time point (Table 2). In the BMP-2 treated defects, cortical bridging also occurred 1 week earlier compared to the 0.9mm defect group from Experiment 1. In contrast, in the collagen group only 1 animal showed cortical bridging and 87.5% manifested as non-unions.

**Table 2.** Number of bridged cortices per callus evaluated weekly in two perpendicular planes and number of mice with successful fracture healing in the collagen and the collagen+BMP-2 groups (≥ 3 bridged cortices, threshold 395 mg HA/cm^3^).

| Week | Group | Number of bridged cortices |  |  |  |  | Fracture healing outcome |  |
| --- | --- | --- | --- | --- | --- | --- | --- | --- |
|  |  | 0 | 1 | 2 | 3 | 4 | Not healed | Healed |
| 0 | collagen | 8 | 0 | 0 | 0 | 0 | 8 | 0 |
|  | collagen+BMP | 8 | 0 | 0 | 0 | 0 | 8 | 0 |
| 1 | collagen | 8 | 0 | 0 | 0 | 0 | 8 | 0 |
|  | collagen+BMP | 8 | 0 | 0 | 0 | 0 | 8 | 0 |
| 2 | collagen | 7 | 0 | 1 | 0 | 0 | 8 | 0 |
|  | collagen+BMP | 0 | 0 | 0 | 0 | 8 | 0 | 8 |
| 3 | collagen | 7 | 0 | 0 | 1 | 0 | 7 | 1 |
|  | collagen+BMP | 0 | 0 | 0 | 0 | 8 | 0 | 8 |
| 4 | collagen | 7 | 0 | 0 | 1 | 0 | 7 | 1 |
|  | collagen+BMP | 0 | 0 | 0 | 0 | 8 | 0 | 8 |
| 5 | collagen | 7 | 0 | 0 | 1 | 0 | 7 | 1 |
|  | collagen+BMP | 0 | 0 | 0 | 0 | 8 | 0 | 8 |
| 6 | collagen | 7 | 0 | 0 | 1 | 0 | 7 | 1 |
|  | collagen+BMP | 0 | 0 | 0 | 0 | 8 | 0 | 8 |

### Histology

As shown by Safranin-O staining six weeks after osteotomy, hardly any cartilage residuals were present in the defect region in all groups, indicating progression of the healing process to the final remodelling stage (Fig. 5, Row 1 and 2). To visualize potential remnants of the collagen scaffolds in Experiment 2, 1-2 samples/group were stained with Sirius-Red (Fig. 5, Row 3 and 4). For comparisons, we also included sections of the original collagen scaffolds (Fig. 5, Row 5), where red color of the filaments is indicative of collagen. In contrast, no red signal indicative of collagen was seen in the defect center of both groups (Fig. 5, Row 4). This suggested complete degradation of the scaffolds with accumulation of fat cells in the defect (unbridged defects in collagen group and in the restored medullary cavity, bridged defects in the collagen+BMP-2 group; Fig. 5, Row 4).

**Figure 5.**
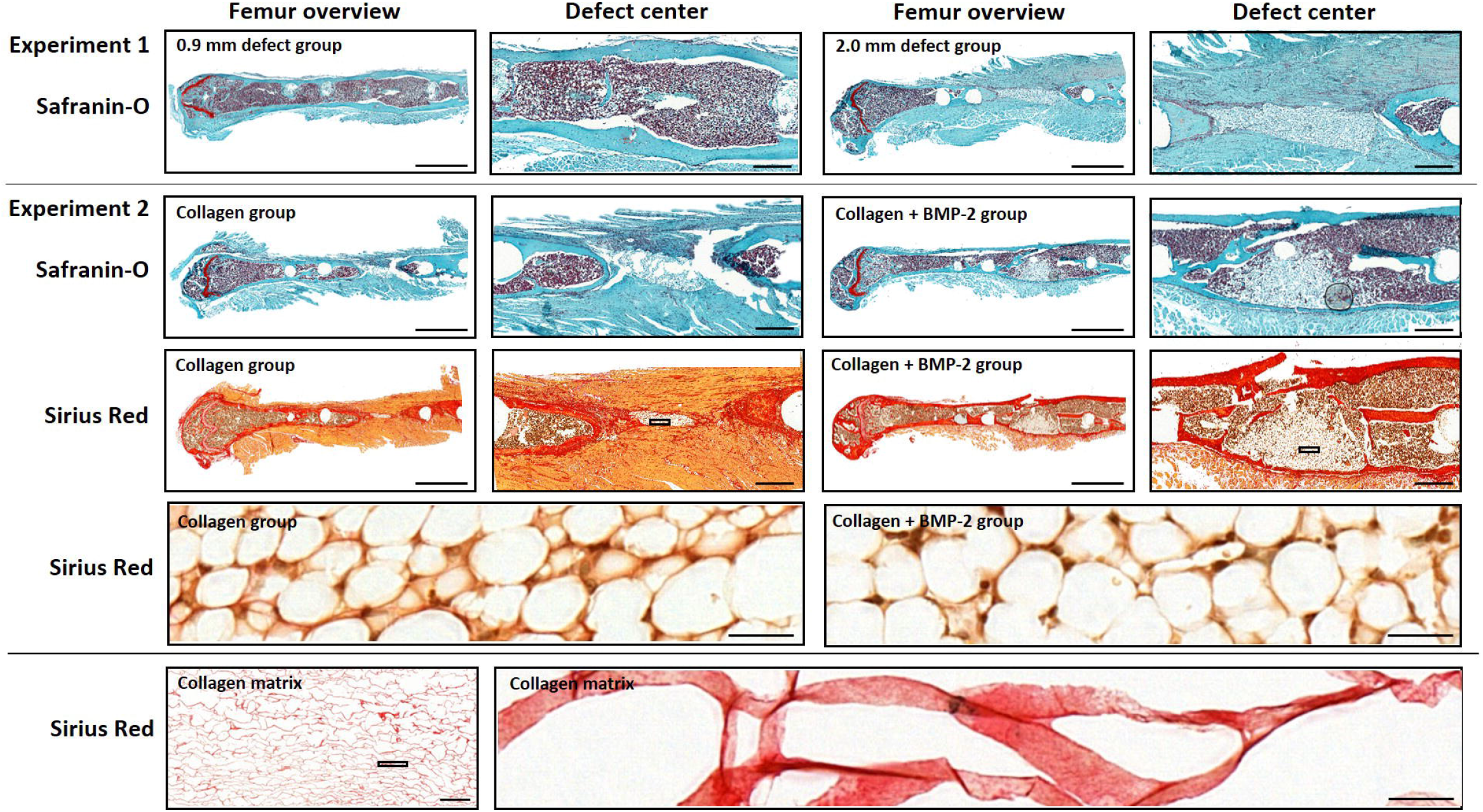
Histology of longitudinal sections of fractured femora 6 weeks after defect surgery. Row 1: Safranin-O staining of 0.9mm and 2.0mm defect groups - overview images, scale bar = 1 mm and area between inner pins of fixator, scale bar = 100 μm. Row 2: Safranin-O staining of collagen and collagen+BMP-2 groups - overview images, scale bar = 1 mm and area between inner pins of fixator, scale bar = 100 μm. Row 3: Sirius-Red staining of collagen and collagen+BMP-2 groups - overview images, scale bar = 1 mm and area centered between inner pins of fixator, scale bar = 100 μm. Row 4: Sirius-Red staining of defect center of collagen group and collagen+BMP-2 group, scale bar = 50 μm. Row 5: Sirius-Red staining of original collagen matrix, scale bar = 500 μm (left image) and 50 μm (right image).

## Discussion

In this study, we showed that time-lapsed *in vivo* micro-CT allows for spatio-temporal discrimination between physiological and impaired fracture healing patterns in a mouse femur defect model. For this purpose, we compared the healing pattern in three callus sub-volumes and the adjacent cortical fragments in a small (0.9mm) and a large (2.0mm) defect group using a previously developed two-threshold density micro-CT protocol ^21,22^. In a second step, we applied the micro-CT based monitoring approach for the characterization of the bone regeneration capacity of functionalized biomaterials using porous collagen scaffolds +/- bone morphogenetic protein (BMP-2).

By registering consecutive scans, we were able to include dynamic parameters such as bone formation and resorption in our micro-CT based monitoring approach as previously described ^21^. In the small defect group (0.9mm), this allowed for characterization of the different healing phases seen by changes in formation and resorption in the osseous callus volume. Specifically, we saw that the initiation of bone formation (maximum in week 1-2), indicating the onset of the reparative phase, triggered bone resorption (maximum in week 2-3) thereby initiating the remodelling phase. Maximum osseous callus volumes were observed in week 3, which remained stable until the study end, while the callus mineralization increased throughout the healing period. The observed bone turnover (bone formation triggering bone resorption) and callus maturation (increasing mineralization of initial callus) patterns are similar to a recent study in 1.5mm defects, which showed an uncompromised healing pattern ^22^. However, the smaller defect size (− 40%) in the current study, changed the temporal occurrence of peaks in bone formation and resorption, which were seen one week earlier compared to the previous study in 1.5mm defects, indicating faster healing progression.

As expected from literature ^23^, the larger defects (2.0mm) showed impaired healing predominantly leading to non-unions in 7 of 8 animals at the study end (Fig. 2, Table 1). This is in line with a study by Zwingenberger at al. ^23^, which previously showed a strong correlation of the defect size and non-unions assessed via *in vivo* X-ray measurements as well as end point micro-CT and histological analyses in a similar defect model. Via registration of consecutive *in vivo* micro-CT scans, we were now able to reveal and visualize the preceding spatio-temporal deviations in callus formation and remodelling compared to the uncompromised healing pattern seen in the 0.9mm defect group. Deviations from normal healing occurred in all callus sub-volumes (defect center, defect periphery, cortical fragment periphery), although initiation of bone formation was not changed spatio-temporally, starting in the same sub-volumes (DC, FP) and at the same time (week 0-1 to week 1-2). However, the rise in bone formation was significantly lower endosteally and the duration of the bone formation was reduced periosteally compared to the 0.9mm defect group. This also led to only small endosteal and peripheral callus dimensions with the periosteal callus volume even decreasing to baseline values by the study end. Under normal healing conditions, callus formation in this femur defect model proceeds from both the endosteal and the periosteal regions to the defect periphery ^22,24^, whereas in large defects hardly any bone formation was seen here, suggesting early termination of the fracture healing process.

It is important to separately look at the adjacent cortical fragments ^22^, where we previously saw an opposite trend in bone turnover compared to the callus sub-volumes. Whereas bone formation was the dominant factor in the callus VOIs, bone resorption was the main factor in the adjacent cortex. In the current study, increased defect size was associated with a significant higher decline in cortical bone volume (−67% vs. −19%) and bone mineral density (−16% vs. −6%; Fig. 1 and Fig. 2). Peri-fracture bone resorption and demineralization of cortical bone have also been previously reported in preclinical and clinical studies and were associated with stiff fixation leading to stress shielding of the bone ^25,26^. Our recently developed time-lapsed *in vivo* micro-CT monitoring approach therefore allowed to precisely characterize the spatio-temporal changes in structural and dynamic callus parameters preceding non-unions. In a next step, we applied this monitoring approach to evaluate the bone regeneration potential of biomaterials using collagen +/- BMP-2 as test materials. We selected collagen as test material due to its wide usage in scaffolds intended for promotion of bone regeneration and the comprehensive literature on its performance in *in vitro*, preclinical and clinical applications ^27–29^. In our study, the application of collagen scaffolds did not significantly improve bone regeneration compared to empty defects of the same size, with the same non-union rates being observed at the study end (87.5% in both groups). Although periosteal bone formation was more pronounced in the collagen group, the fracture healing outcome was not different between groups. This is in line with previous bone defect studies in different rodent models (loaded and non-loaded bones) reporting no significant effect of collagen scaffolds on bone regeneration with incomplete bone union or non-unions at the study end ^30,31^. Similarly, in large animal models collagen scaffolds either failed to prevent non-unions ^32^ or to restore the mechanical bone properties ^33^. The low regeneration potential of collagen scaffolds has been attributed to a general lack of bioactivity of collagen despite its overall favorable features such as low antigenicity, high biodegradability and high biocompatibility ^27–29^. Furthermore, studies indicate a high relevance of the pore size in the collagen scaffolds, with smaller pores potentially being affected by cellular occlusion, and preventing cellular penetration, production of extracellular matrix and neovascularization of the inner areas of the scaffold ^28^. In contrast, larger pores have been associated with low mechanical scaffold properties and early scaffold degradation ^34,35^. A major limitation of most studies is that they did not include empty defect and positive controls, which would be essential for better understanding of specific changes in healing patterns mediated by different biomaterials.

To further promote bone healing, scaffolds have been incubated in solutions containing growth factors (e.g. BMP-2, BMP-7; ^36,37^). Especially BMP-2 has been widely applied in preclinical studies to promote healing of critical sized defects (for review see ^38^) and in clinical settings. However, studies reported BMP-2 associated adverse events and complications (e.g. antibody formation, inflammation, ectopic bone formation, carcinogenicity ^39–41^). Therefore, using low BMP-2 dosages seems crucial in order to avoid adverse effects while preserving its osteoanabolic potential ^42^. So far preclinical studies used a wide range of BMP-2 dosages (0.1-150.000 μg BMP-2 per defect in small animals as summarized in a comprehensive review by Gothard et al. ^38^). In this study, we therefore down-scaled the BMP-2 dosages from previous studies (low BMP-2 dosages with beneficial effect on bone healing; 10/75 μg BMP-2 per defect ^43,44^) to the volume of our scaffold (2 μg BMP-2 per defect). Similar to other studies ^38^, BMP-2 application significantly accelerated the healing process leading to early cortical bridging. However, when looking at the defect sub-volumes separately, we saw that the spatio-temporal healing pattern largely differed from the physiological healing pattern seen in the 0.9mm group. In the collagen+BMP-2 group the increase in the bone formation rate was higher but persisted for a shorter amount of time compared to the 0.9mm defect group. Furthermore, bone formation mainly took place in the periosteal regions and the defect periphery, whereas in the 0.9mm defect group bone formation was predominantly seen in the defect center. This supports the assumption that the collagen scaffold itself might have hindered cells from migrating into the defect center. In line with other studies ^30,45^, the collagen scaffolds were degraded in both groups by the end of the study as shown by absence of collagen fibers in the defect center via Sirius Red staining of histological sections. Accumulation of fat cells inside the formed cortical shell (unions) or in the defect center (non-unions) indicated differentiation of mesenchymal stem cells (MSCs) towards the adipogenic lineage in the later healing period. Concluding from the time-lapsed *in vivo* micro-CT evaluation, strong BMP-2 induced bone formation associated with osteogenic differentiation of MSCs was only transiently observed in the early healing period. Removal of the BMP-2 induced stimulus in later healing phases might have caused a change from osteogenic to adipogenic differentiation of MSCs. This shift might have been further strengthened by stress shielding of the defect region associated with stiff external fixation ^46^.

Our study therefore shows that sub-volume specific characterization of functionalized biomaterials seems crucial for assessing their bone regeneration potential. The study further indicates that time-lapsed *in vivo* micro-CT combined with our recently developed two-threshold density approach should be used in future studies to identify parameters for prediction of non-unions in early healing phases. Via registration of consecutive scans as described recently, we were able to precisely characterize and understand which spatio-temporal deviations led to non-union formation and how this was prevented by BMP-2 application. Using collagen and BMP-2 as test materials, the time-lapsed *in vivo* micro-CT-based monitoring approach was shown to be suitable for spatio-temporal assessment of callus formation and remodelling patterns and could be used in future studies to characterize and precisely understand the regeneration capacity of functionalized biomaterials.

The current study has several limitations. We did not track the degradation of the collagen scaffolds over time and only performed end-point histological assessment of collagen residuals via Sirius Red staining. Future studies should apply fluorescent biomaterials and reporters, which could be visualized via *in vivo* optical imaging ^47–49^. We did also not measure the BMP-2 release kinetics. This could be similarly achieved via the use of labeled growth factors^50^ in combination with multimodal imaging (e.g. optical imaging, PET, SPECT, micro-CT)^17,51^. Furthermore, in order to reliably create critical-sized defects, the defect size would need to be increased by ca. 25% in future studies.

Nevertheless, application of time-lapsed *in vivo* micro-CT allows faster and more precise evaluation of biomaterials for bone regeneration, thereby also reducing the animal numbers involved according to the 3R’s of animal welfare. It could further be used for early prediction of non-union formation and identification of biomarkers. The approach could be supplemented with the registration of 2D-histology section into the 3D-micro-CT volume^52^, potentially allowing for a spatio-temporal understanding of molecular and cellular changes induced by different biomaterials. To better mimic the clinical situation, future studies should also assess the performance of biomaterials under load application, which could be achieved using a recently developed loading fixator^53^. By combining our *in vivo* time-lapsed micro-CT based monitoring approach with individualized loading regimes, this would allow for thorough characterization of biomaterials under clinically relevant loading conditions.

To conclude, *in vivo* time-lapsed micro-CT allows (1) spatio-temporal discrimination between normal and disturbed healing patterns relevant for the detection of distinct features associated with different medical conditions and (2) spatio-temporal characterization of the bone regeneration capacity of functionalized biomaterials.

## Materials and Methods

### Animals

All animal procedures were approved by the local authorities (license number: 36/2014; Kantonales Veterinäramt Zürich, Zurich, Switzerland). We confirm that all methods were carried out in accordance with relevant guidelines and regulations (Swiss Animal Welfare Act and Ordinance (TSchG, TSchV)) and reported considering ARRIVE guidelines. Female 12 week-old C57BL/6J mice were purchased from Janvier (Saint Berthevin Cedex, France) and housed in the animal facility of the ETH Phenomics Center (EPIC; 12h:12h light-dark cycle, maintenance feed (3437, KLIBA NAFAG, Kaiseraugst, Switzerland), 5 animals/cage). At an age of 20 weeks, all animals received a femur defect. In Experiment 1, one group of animals received a small defect (defect length: 0.86±0.09mm, n=10) and a second group received a large defect (defect length: 2.00±0.19mm, n=8; housing after surgery: 2-3 animals/cage). In Experiment 2 both groups received a 2mm femur defect with application of either collagen scaffolds (d=2mm, h=2mm; ILS, Saint Priest, France; n=8) or collagen scaffolds+BMP-2 (2.5μg/scaffold; PeproTech, London, UK; n=8). Perioperative analgesia (25 mg/L, Tramal^®^, Gruenenthal GmbH, Aachen, Germany) was provided via the drinking water two days before surgery until the third postoperative day. For surgery and micro-CT scans, animals were anesthetized with isoflurane (induction/maintenance: 5%/1-2% isoflurane/oxygen).

### Femur defect

An external fixator (Mouse ExFix, RISystem, Davos, Switzerland; stiffness: 24N/mm^21^) was positioned at the craniolateral aspect of the right femur and attached using four mounting pins. First, the most distal pin was inserted approximately 2mm proximal to the growth plate, followed by placement of the most proximal and the inner pins. Subsequently, a femur defect was created using 1 and 2 Gigli wire saws for the small and the large defect, respectively.

### Time-lapsed *in vivo* micro-CT

Immediate post-surgery correct positioning of the fixator and the defect was visualized using a vivaCT 40 (Scanco Medical AG, Brüttisellen, Switzerland) (isotropic nominal resolution: 10.5 μm; 55 kVp, 145 μA, 350 ms integration time, 500 projections per 180°, 21 mm field of view (FOV), scan duration ca. 15 min). Subsequently, the defect region as well as the adjacent cortex were scanned weekly (week 1-6) with the same settings and morphometric indices and mineralization progression were determined in four volumes of interest (for details on methods see ^21,22^): defect center (DC), defect periphery (DP), cortical fragment center (FC), and fragment periphery (FP). Data were normalized to the central VOIs: DC/DC, DP/DC, FC/FC, FP/FC. Cortical bridging was assessed as previously described in ^22^.

### Animations

To further visualize the defect healing process and the VOIs involved, 3-dimensional computer renderings were created from the time-lapsed micro-CT data for one mouse in each respective group across both experiments (0.9mm defect group, 2.0mm defect group, collagen scaffold, collagen scaffold+BMP-2), then animated and captioned to create videos (Supplementary Video 1, Supplementary Video 2). Bone mineral density data was single-value thresholded (BMD = 395 mg HA/cm^3^) at each time point to create a binary array, simplifying the rendering process. This binarized bone location data was interpolated using a Euclidean Distance Transform, generating data for time points in between each weekly measurement. This data was used to create smooth, animated transitions from one measurement to the next, for visualization purposes, and was not used in any analysis. To indicate bone formation and resorption between each time point, bone remodelling data from the time-lapsed *in vivo* micro-CT method was taken ^21^. Depictions of the VOIs were rendered directly from their location data used in this approach. All renderings of the 3D data were performed in ParaView (Kitware, Version 5.6; Clifton Park, NY). All numerical analysis used for the animations was done using custom Python 3 scripts. Video editing and captioning was done in Hitfilm Express (FXhome, Version 13; Norwich, UK).

### Histology

To exemplarily visualize the defect region, histology was performed in 1-2 animals/group. On day 42 femurs were excised, the femoral head was removed, and the samples were placed in 4% neutrally buffered formalin for 24 hours and subsequently decalcified in 12.5% EDTA for 10-14 days. The samples were embedded in paraffin and 4.5 μm longitudinal sections were stained with Safranin-O/Fast Green: Weigert’s iron haematoxylin solution (HT1079, Sigma-Aldrich, St. Louis, MO) - 4min, 1:10 HCl-acidified 70% ethanol – 10s, tap water - 5min, 0.02% Fast Green (F7258, Sigma-Aldrich, St. Louis, MO) - 3min, 1% acetic acid - 10s, 0.1% Safranin-O (84120, Fluka, St. Louis, MO) - 5min. In order to visualize remnants of the collagen scaffolds, Sirius Red staining was performed in 1 animal from the collagen and 1 animal from the collagen+BMP-2 group. For comparison, we also stained sections of the original collagen matrix with Sirius-Red: Weigert’s iron haematoxylin solution (HT1079, Sigma-Aldrich, St. Louis, MO) - 8min, tap water - 10min, 1:9 Picro-Sirius Red solution (Picric acid: 80456, Fluka, St. Louis, MO; Direct Red 80: AB133584, ABCR, Karlsruhe, Germany) - 1h, 5% acetic acid 2×10s. For both stainings, images were taken with Slide Scanner Pannoramic 250 (3D Histech, Budapest, Hungary) at 20x magnification.

### Statistics

CT analysis: Data were tested for normal distribution (Shapiro-Wilk-Test) and homogeneity of variance (Levene-Test). Depending on the test outcome, group comparisons of data derived at single time points were done by Student’s t-test or Mann-Whitney U-tests (IBM SPSS Statistics Version 23). For statistical evaluation of repeated measurements two-way ANOVA with Geisser-Greenhouse correction and Bonferroni correction (GraphPad Prism 8) were performed. The level of significance was set at *p* < 0.05.

## Author Contributions Statement

The study was designed by E.W., G.A.K., S.H. and R.M.. The *in vivo* experiments were performed by E.W., G.A.K. and E.F.. Histological stainings were performed by E.W. and M.H.N.. Data analyses were performed by E.W. and D.C.T.B.. Illustrations of VOIs and animations on callus formation and remodelling were made by B.J.S.. The manuscript was written by E.W. and reviewed and approved by all authors.

## Data availability

All necessary data generated or analyzed during the present study are included in this published article and its Supplementary Information files (preprint available on BioRxiv (BIORXIV/2020.10.02.324061). Additional information related to this paper may be requested from the authors.

## Acknowledgements

The authors gratefully acknowledge support from the EU (BIODESIGN FP7-NMP-2012-262948 and ERC Advanced MechAGE ERC-2016-ADG-741883). E. Wehrle received funding from the ETH Postdoctoral Fellowship Program (MSCA-COFUND, FEL-25_15-1).

