## Supplementary Information for "Longitudinal *in vivo* micro-CT-based approach allows spatio-temporal characterization of fracture healing patterns and assessment of biomaterials in mouse femur defect models"

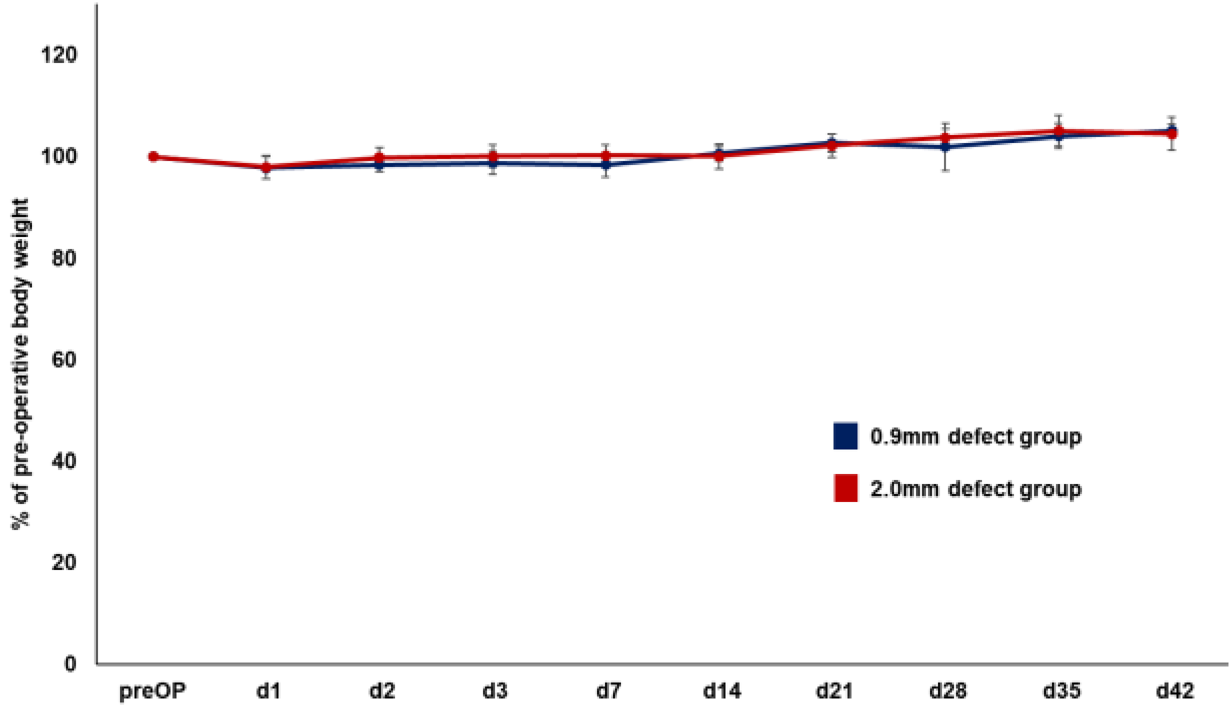


**Supplementary Fig. S1.** *In vivo* monitoring of body weight of the mice from the 0.9mm defect group (n=8) and the 2.0mm defect group (n=10) measured pre-operatively (preOP), on postoperative days 1-3 and weekly from day 7 to day 42. The postoperative values were related to the preoperative data.

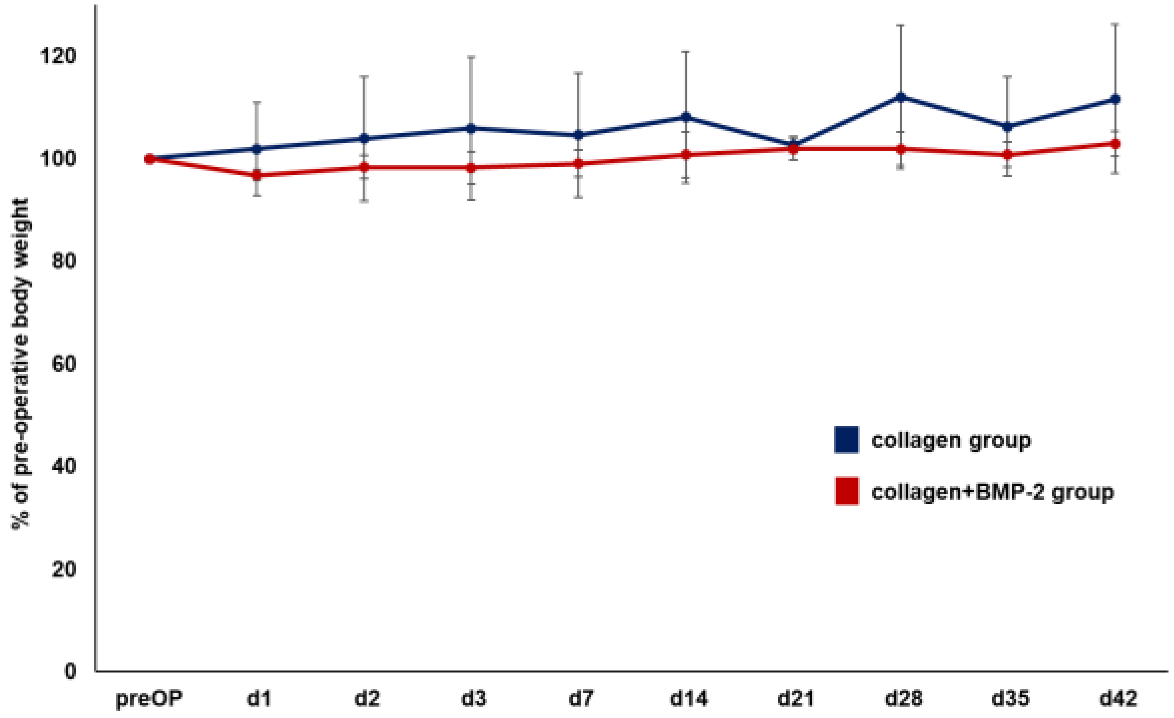


**Supplementary Fig. S2.** *In vivo* monitoring of body weight of the mice from the collagen group (n=8) and the collagen+BMP-2 group (n=8) measured pre-operatively (preOP), on postoperative days 1-3 and weekly from day 7 to day 42. The postoperative values were related to the preoperative data.

**Supplementary Table S1.** Study design (female 20 week-old C57BL/6J mice)

| Experiment | Group | Group size | Size of femur defect | Biomaterial application | *In vivo* micro-CT measurements | Registration of micro-CT scans # | Histology |
|---|---|---|---|---|---|---|---|
| 1 | 0.9mm | n=11 | 0.9mm (n=10) | - | d0, week 1-6 (n=10) | week 1-6 to week 0-5 (n=10) | week 6 (n=2) |
| | 2.0mm | n=8 | 2.0mm (n=8) | - | d0, week 1-6 (n=8) | week 1-6 to week 0-5 (n=7) | week 6 (n=2) |
| 2 | collagen | n=8 | 2mm (n=8) | collagen | d0, week 5+6 (n=8) | week 1-6 to week 0-5 (n=8) | week 6 (n=1) |
| | collagen +BMP-2 | n=8 | 2mm (n=8) | collagen +BMP-2 | d0, week 5+6 (n=8) | week 1-6 to week 0-5 (n=8) | week 6 (n=1) |

### micro-CT scan taken at timepoint x registered to micro-CT scan taken at timepoint x-1

**Supplementary Video 1.** Visualisation of the defect healing process and the VOIs involved for a representative animal from the 0.9mm defect group and the 2.0mm defect group.

**Supplementary Video 2.** Visualisation of the defect healing process and the VOIs involved for a representative animal from the collagen group and the collagen+BMP-2 group).
